# Turing’s diffusive threshold in random reaction-diffusion systems

**DOI:** 10.1101/2020.11.09.374934

**Authors:** Pierre A. Haas, Raymond E. Goldstein

**Affiliations:** Mathematical Institute, University of Oxford, Woodstock Road, Oxford OX2 6GG, United Kingdom; Department of Applied Mathematics and Theoretical Physics, Centre for Mathematical Sciences, University of Cambridge, Wilberforce Road, Cambridge CB3 0WA, United Kingdom

## Abstract

Turing instabilities of reaction-diffusion systems can only arise if the diffusivities of the chemical species are sufficiently different. This threshold is unphysical in most systems with *N* = 2 diffusing species, forcing experimental realizations of the instability to rely on fluctuations or additional nondiffusing species. Here we ask whether this diffusive threshold lowers for *N* > 2 to allow “true” Turing instabilities. Inspired by May’s analysis of the stability of random ecological communities, we analyze the probability distribution of the diffusive threshold in reaction-diffusion systems defined by random matrices describing linearized dynamics near a homogeneous fixed point. In the numerically tractable cases *N* ≤ 6, we find that the diffusive threshold becomes more likely to be smaller and physical as *N* increases and that most of these many-species instabilities cannot be described by reduced models with fewer species.

---

In 1952, Turing described the pattern-forming instability that now bears his name [1]: diffusion can destabilize a fixed point of a system of reactions that is stable in well-mixed conditions. Nigh on threescore and ten years on, the contribution of Turing’s mechanism to chemical and biological morphogenesis remains debated, not least because of the *diffusive threshold* inherent in the mechanism: chemical species in reaction systems are expected to have roughly equal diffusivities, yet Turing instabilities cannot arise at equal diffusivities [2, 3]. It remains an open problem to determine how much of a diffusivity difference is required for generic systems to undergo this instability, yet this diffusive threshold has been recognized at least since reduced models of the Belousov–Zhabotinsky reaction [4, 5] only produced Turing patterns at unphysically large diffusivity differences.

For this reason, the first experimental realizations of Turing instabilities [6–8] obviated the threshold by using gel reactors that greatly reduced the effective diffusivity of one species [9, 10]. (Analogously, biological membrane transport dynamics can increase the effective diffusivity difference [11].) Later work showed how binding to an immobile substrate, or more generally, a third, nondiffusing species, can allow Turing instabilities even if the *N* = 2 diffusing species have equal diffusivities [12–14]. Such nondiffusing species continue to permeate more recent work on the network topology of Turing systems [15, 16].

Moreover, Turing instabilities need not be deterministic: fluctuation-driven instabilities in reaction-diffusion systems have noise-amplifying properties that allow their pattern amplitude to be comparable to that of deterministic Turing patterns [17], with a lower diffusive threshold than the deterministic one [18-21]. A synthetic bacterial population with *N* = 2 species that exhibits patterns in agreement with such a stochastic Turing instability, but does not satisfy the conditions for a deterministic instability [22], was reported recently.

These experimental instabilities relying on fluctuations or the dynamics of additional nondiffusing species and the nonlinear instabilities arising from finite-amplitude perturbations [2] are not however instabilities in Turing’s own image. Can such instabilities be realized instead in systems with *N* > 2 diffusing species? Equivalently, is the diffusive threshold lower in such systems? These questions have remained unanswered, perhaps because, in contrast to the textbook case *N* = 2 and the concomitant picture of an “inhibitor” out-diffusing an “activator” [23, 24], the complicated instability conditions for *N* > 2 [25] do not lend themselves to analytical progress.

Here, we analyze the diffusive threshold for Turing instabilities with 2 ≤ *N* ≤ 6 diffusing species. Inspired by May’s work on the stability of random ecological communities [26], we analyze *random Turing instabilities* by sampling random matrices that represent the linearized reaction dynamics of otherwise unspecified reaction-diffusion systems. A semianalytic approach shows that the diffusive threshold is more likely to be smaller and physical for *N* = 3 compared to *N* = 2, and that two of the three diffusivities are equal at the transition to instability. We extend these results to the remaining numerically tractable cases of reaction-diffusion systems with 4 ≤ *N* ≤ 6 and two different diffusivities: their Turing instabilities are still more likely to have a smaller and physical diffusive threshold, but most of them cannot be described by reduced models with fewer species.

We begin with the simplest case, *N* = 2, in which species *u, v* obey

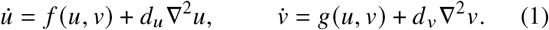

The conditions for Turing instability in this system [24] only depend on the four entries of the Jacobian

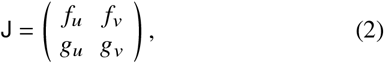

the partial derivatives of the reaction system at a fixed point (*u*_*_,*v*_*_) of the homogeneous system. This fixed point is stable to homogeneous perturbations iff *J* ≡ det J > 0 and *I*_1_ ≡ tr J < 0. A stable fixed point of this kind is unstable to a Turing instability only if *p* ≡ – *f_u_ g_v_* > 0 [24]. Defining the diffusion coefficient ratio *D*_2_ = max {*d_u_/d_v_, d_v_*/*d_u_* } ≥ 1, a Turing instability occurs iff these conditions hold along with [27]

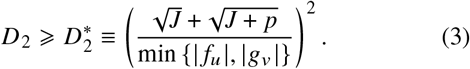

This diffusivity difference 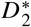, which is required mathematically for instability, is unphysical [Fig. 1(a)] if it exceeds the diffusivity difference 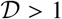 of the physical system: 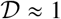 for similarly sized molecules in solution, but, e.g., 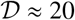 for the stochastic Turing instability observed in Ref. [22]. Hereinafter we take 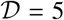 arbitrarily (but have checked that the value of 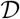 does not affect results qualitatively). To quantify 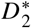, we introduce the *range R* of kinetic parameters,

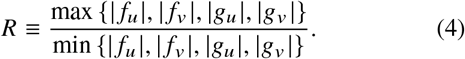

**FIG. 1.**
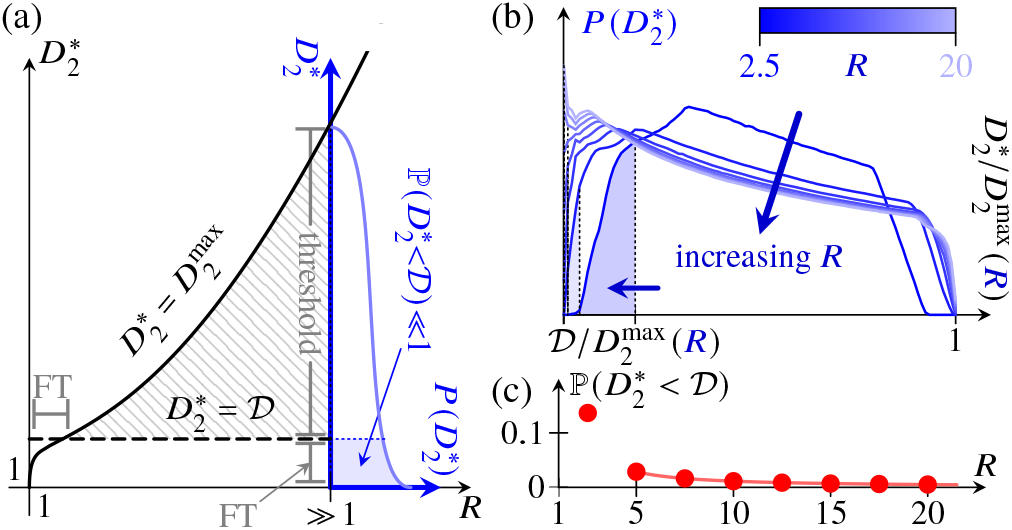
Turing’s diffusive threshold for *N* = 2. (a) Cartoon of the diffusive threshold and the fine-tuning (FT) problem for *R* ≈ 1 and *R* » 1. The diffusivity difference required mathematically is unphysical in the hatched region 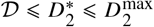. (b) Distribution 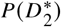, supported on the (scaled) interval 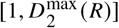, estimated for different *R*. (c) Plot of 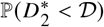 [shaded areas in panels (a) and (b)] against *R*, revealing the diffusive threshold. Markers: estimates from panel (b); solid line: exact result [27] for 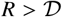 [33].

Equivalently, *f_u_,f_v_,g_u_,g_v_* ∈ *I* ≡ [-*R*,-1] ∪ [1,*R*] up to scaling, with one parameter equal to ±1 and one equal to ±*R*. One deduces [27] that

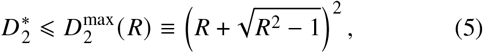

as shown in Fig. 1(a). Note that 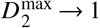 as *R* → 1; in this limit, there is no diffusive threshold: *R* ≈ 1 is a particular instance of the converse *fine-tuning problem* for the reaction kinetics that allows Turing instabilities at nearly equal diffusivities more generally [3]. If *R* » 1, then 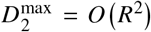. This does not imply the existence of a threshold, for it does not preclude most systems with range *R* having 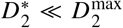. The existence of a diffusive threshold therefore relates to the distribution of 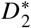 for systems with range *R*.

To understand this distribution, we draw inspiration from May’s statistical analysis of the stability of ecological communities [26], which studies random Jacobians, corresponding to equilibria of otherwise unspecified population dynamics. By analogy, we study random Turing instabilities, sampling uniformly and independently random Jacobians corresponding to equilibria of otherwise unspecified reaction kinetics, and analyze the criteria for them to be Turing unstable. There is of course no more reason to expect the kinetic parameters to be independent or uniformly distributed than there is reason to expect the linearized population dynamics in May’s analysis [26] to be independent or normally distributed. Yet, in the absence of experimental understanding of what the distributions of these parameters should be (in either context), the potential of the random matrix approach to reveal stability principles has been amply demonstrated in population dynamics [34–44].

We sample the kinetic parameters in Eq. (2) independently and uniformly from *I*, set one of them equal to ±1 and one equal to ±*R*, and thus estimate the probability distribution 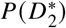 for fixed *R* [Fig. 1(b)]. The threshold is quantified by the probability of a Turing instability being physical,

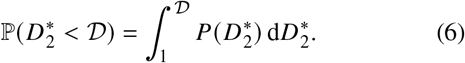

Both from the estimates in Fig. 1(b) and by evaluating the integral in closed form [27], we find that 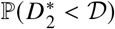 is tiny [Fig. 1(c)], except if *R* is small, which is the fine-tuning problem again. In other words, the required diffusivity difference is very likely to be unphysical. This expresses Turing’s diffusive threshold for *N* = 2.

To investigate how this threshold changes with *N* we consider next the *N* = 3 system

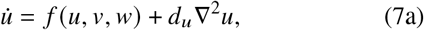

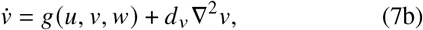

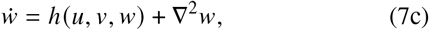

where we have rescaled space to set *d_w_* = 1. We introduce the matrix of diffusivities and the reaction Jacobian,

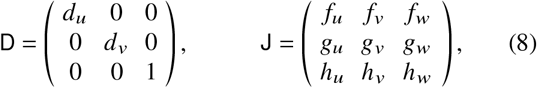

in which the entries of J are again the partial derivatives evaluated at a fixed point (*u*_*_,*v*_*_,*w*_*_) of the homogeneous system. This fixed point is unstable to a Turing instability if it is stable but, for some eigenvalue – *k*^2^ < 0 of the Laplacian, 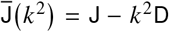 is unstable [3], i.e. has an eigenvalue *λ* such that Re(*λ*) < 0. More precisely, a Turing instability arises when a real eigenvalue of 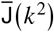 crosses zero, i.e. when 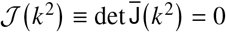, and therefore arises first at a wavenumber *k* = *k*, with 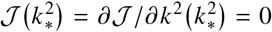 [3]. Hence 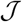, a cubic polynomial in *k*^2^, has a double root at 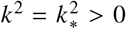, so its discriminant [29] vanishes. This discriminant, Δ(*d_u_,d_v_*), is a polynomial in *d_u_, d_v_*. We denote by *K* (*d_u_, d_v_*) the double root of 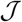 corresponding to a point (*d_u_, d_v_*) on the curve Δ(*d_u_, d_v_*) = 0.

Determining the diffusive threshold for Turing instability in Eqs. (7) thus requires solving the problem

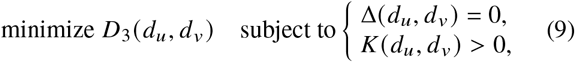

in which the diffusion coefficient ratio is

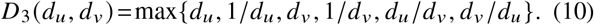

With the aim in mind of obtaining statistics for the minimal value 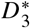, direct numerical solution of this constrained optimization problem is obviously not a feasible approach. In the Supplemental Material [27], we therefore show how to reduce solving problem (9) to polynomial root finding. This semianalytic approach reveals a particular class of minima, attained at the vertices of the contours of *D*_3_ (*d_u_, d_v_*) [Fig. 2(a)], i.e. at *d_u_* = 1, *d_v_* = 1, or *d_u_ = d_v_*. In these cases, Δ(*d_u_, d_v_*) = 0 is a (sextic) polynomial in the single variable *d_v_, d_u_*, or *d* = *d_u_* = *d_v_*, respectively. We call these minima “binary”, since the corresponding systems have only two different diffusivities. We implement this approach numerically [27], and sample random systems similarly to the case *N* = 2, drawing the entries of J in Eq. (8) uniformly and independently at fixed range *R*.

**FIG. 2.**
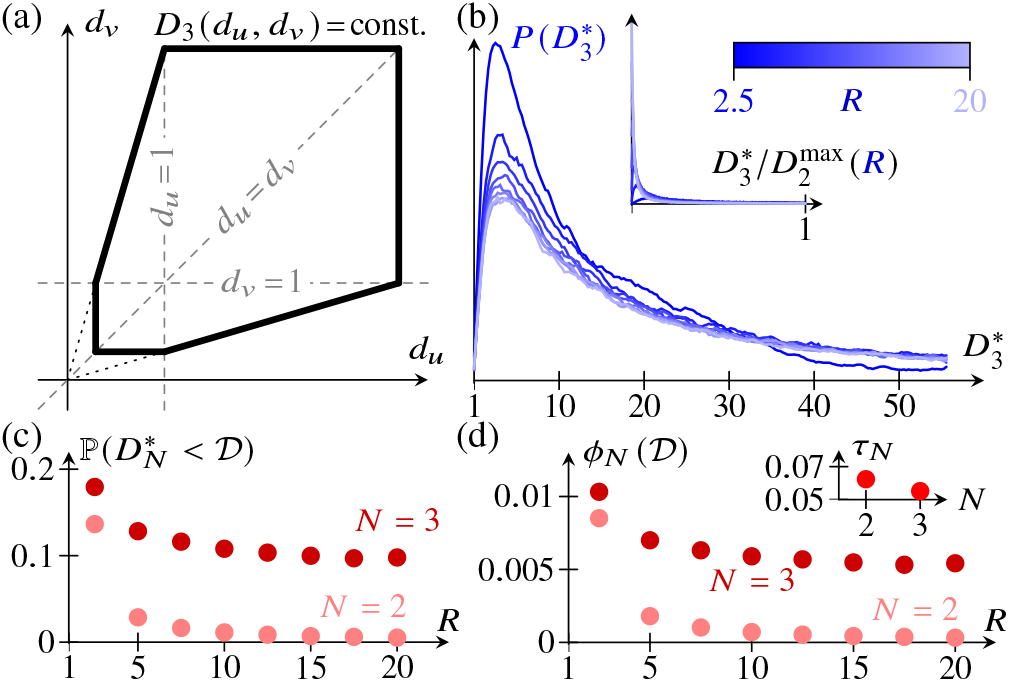
Results for *N* = 3. (a) Contours of *D*_3_ (*d_u_, d_v_*) in the positive (*d_u_, d_v_*) quadrant. (b) Smoothed distribution 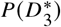, estimated for different *R*. Inset: same plot, scaled to 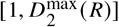 for comparison to *N* = 2 in Fig. 1(a). (c) 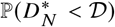 against *R* for *N* ∈ {2,3}: the diffusive threshold lowers for *N* = 3 compared to *N* = 2. (d) Proportion 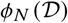 of random Jacobians that have a physical Turing instability, plotted against *R*, for *N* ∈ {2,3}. Inset: proportion of random Jacobians that have a (physical or unphysical) Turing instability, averaged over *R*, for *N* ∈ {2,3} [33].

Remarkably, all global minima we found numerically were binary [27]. This means that the minimizing systems come in two flavors: those with two “fast” diffusers and one “slow” diffuser, and those with one “fast” diffuser and two “slow” diffusers. Systems with a nondiffusing species are a limit of the former; this point will be discussed below. The latter arise in models of scale pattern formation in fish and lizards [45,46], in which short-range pigments respectively activate and inhibit a long-range factor.

The distribution of 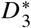, shown for different values of *R* in Fig. 2(b), has a different shape from that of 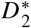 [Figs. 1(a) and 2(b), inset]. While the support of the distribution of 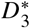 does not appear to be bounded, Fig. 2(c) shows that 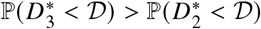. Hence the diffusivity difference is more likely to be physical for *N* = 3 than for *N* = 2: the diffusive threshold is lowered.

The proportion *τ_N_* of random kinetic Jacobians that have a Turing instability (be it physical or unphysical) is smaller for *N* = 3 than for *N* = 2 [Fig. 2(d), inset]. This is not surprising, because a random Jacobian is less likely to correspond to a stable fixed point (which, we recall, is a necessary condition for Turing instability) for *N* = 3 than for *N* = 2, essentially because its entries have to satisfy more conditions for stability if *N* = 3. It is therefore striking that the threshold is reduced sufficiently for *N* = 3 compared to *N* = 2 for the proportion 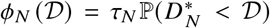 of random Jacobians that have a physical Turing instability to be larger for *N* = 3 than for *N* = 2 [Fig. 2(d)], even though a Turing instability of any kind is more likely if *N* = 2.

To extend these results to *N* > 3 diffusing species, we consider the (linearized) reaction-diffusion system

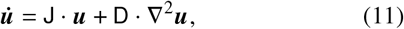

where J is a random kinetic Jacobian, and D is a diagonal matrix of diffusivities. Even with our semianalytic approach, this cannot be analyzed for general D: not even for *N* = 4 were we able to obtain closed forms of the required polynomials. To make further progress, we therefore restrict to binary D in which the *N* diffusivities take two different values only, since we showed above that 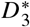 is attained for such binary D. As in the case *N* = 3, this reduces the discriminant condition Δ(D) = 0 to polynomial equations in one variable that determine the minimum diffusivity difference 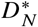 for Turing instability in these binary systems [27].

Figure 3(a) shows that the diffusive threshold lowers further for 4 ≤ *N* ≤ 6 in these systems. At the same time, the fact that most stable random kinetic Jacobians undergo such a binary Turing instability [Fig. 3(b)] suggests that these provide a useful picture of the diffusive threshold. However, *τ_N_* decreases further for *N* > 4 [Fig. 3(c), inset], and the widening of the bottleneck is not sufficient to prevent 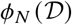 from decreasing for *N* ≤ 4. Nonetheless, since both 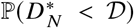 and the proportion of stable random Jacobians that are Turing unstable increase [Figs. 3(a) and 3(b)], so does the proportion of stable random Jacobians that have a physical Turing instability.

**FIG. 3.**
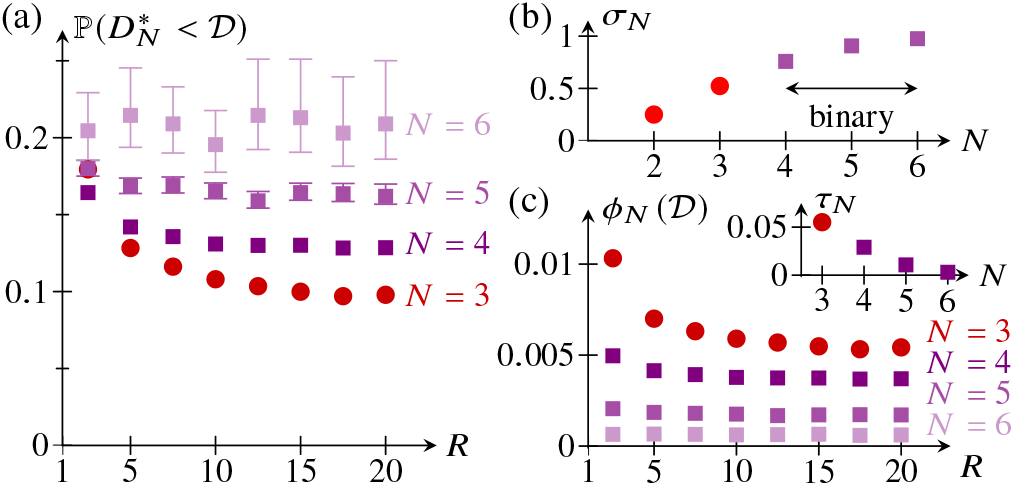
Results for “binary” systems with 4 ≤ *N* ≤ 6. (a) 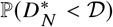 against *R* for 3 ≤ *N* ≤ 6, revealing further lowering of the diffusive threshold compared to the case *N* = 3. (b) Proportion *in* of random stable kinetic Jacobian that have a (binary, if *N* > 3) Turing instability, averaged over *R*, and plotted against *N*. (c) Proportion 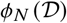 of random Jacobians that have a physical Turing instability plotted against *R*, for 3 ≤ *N* ≤ 6. Inset: proportion *τ_N_* of random Jacobians that have a (physical or unphysical) Turing instability, averaged over *R*, for 3 ≤ *N* ≤ 6 [33].

How then to realize “true” Turing instabilities experimentally? Our analysis shows that the diffusive threshold of a Turing instability is more likely to be physical the more species there are, but how to find an experimental Turing instability in the first place? Turing instabilities remain rare in random reaction systems even as the number of species is increased, but the above shows that this rareness mainly results from the rareness of stable equilibria in such systems. The proverbial search for the needle in a haystack can therefore be avoided by exploring biochemical systems that admit a stable equilibrium, and evolving them towards a “true” Turing instability.

This analysis does not however reveal whether these instabilities lead to patterns that are observable at the physical scale of the system. Analysis of the wavenumber at which the linear instability first arises [27] suggests that we can extend our conclusions: Turing instabilities with more species are more likely to have physical diffusivity differences and to be observable. However, our statistical, linearized analysis cannot fully answer this question of observability, because it fundamentally depends on the system through details of the nonlinearities of its reaction kinetics, which set the precise nature and scale of the Turing patterns that develop beyond onset of the instability; this is why we have relegated this discussion to the Supplemental Material [27].

The different species in the systems with 3 ≤ *N* ≤ 6 analyzed above separate into “fast” and “slow” diffusers. The diffusion of these “slow” species is often ignored in the analysis of systems of many chemical reactions [30], such as the full Belousov-Zhabotinsky reaction [47]. Corresponding reduced models are obtained by substituting the steady-state kinetics of the “slow” species into the remaining equations, thereby eliminating them from the system [30]. The conditions for Turing instability in these reduced models are (almost) equivalent to those for the full model with nondiffusing “slow” species [30]. However, the diffusion of the “slow” species cannot in general be ignored: up to reordering species and rescaling space,

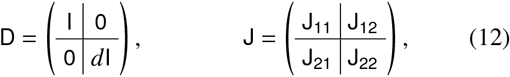

where *d* < 1 is the common diffusivity of the slow diffusers. Results of Ref. [30] imply that there is a Turing instability with nondiffusing “slow” species, i.e. with *d* = 0, only if 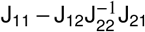 has a positive (real) eigenvalue [27]. Although the proportion of Turing unstable systems that have *n* ≥ 2 fast diffusers (and hence could *a priori* still undergo a Turing instability with *d* = 0) is large [Fig. 4(a)], the proportion of systems that do undergo such an instability is small, even if we restrict to those systems with physical diffusivity differences [Fig. 4(b)]. Hence most of these Turing instabilities with *N* > 2 species require all species to diffuse.

**FIG. 4.**
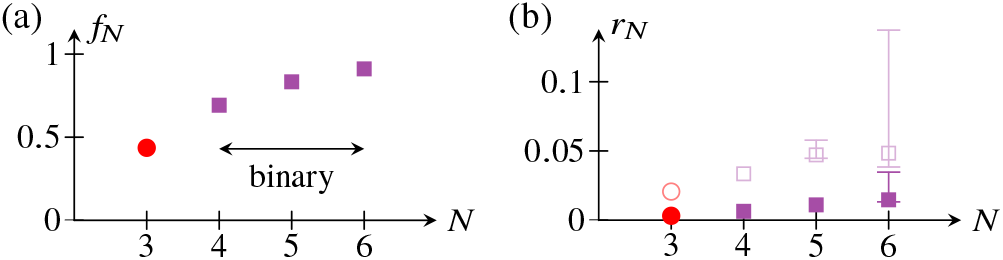
“Slow” diffusers in binary Turing instabilities with 3 ≤ *N* ≤ 6. (a) Proportion *f_N_* of Turing unstable systems with *n* ≥ 2 “fast” diffusers plotted against *N*, averaged over *R*. (b) Proportion of systems that remain Turing unstable at *d* = 0, plotted against *N*, averaged over *R*. Closed markers: all Turing systems; open markers: physical Turing systems with 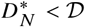 [33].

These many-species Turing instabilities, although binary, are thus more general than the instabilities of systems with nondiffusing species realized experimentally in gel reactors [6–8] and analyzed theoretically in Ref. [30]. In particular, this shows that reduced models give but an incomplete picture of Turing instabilities. Together with our main result, that the diffusive threshold lowers as *N* increases, this implies that the failure of a reduced model to produce a physical Turing instability cannot be taken as an indication that a Turing instability cannot exist in the full system that the reduced model seeks to describe.

In this Letter, we have analyzed random Turing instabilities to show how the diffusive threshold that has hampered experimental efforts to generate “true” Turing instabilities in systems of *N* = 2 diffusing species lowers for systems with *N* ≥ 3, most of whose instabilities cannot be described by reduced models with fewer species. All of this does not, however, explain the existence of a “large” threshold in the first place: even though Turing instabilities at equal diffusivities are impossible [2, 3], this does not mean that the threshold needs to be “large”. In this context, we prove an asymptotic result in the Supplemental Material [27]: for a Jacobian J to allow a Turing instability at almost equal diffusivities D ≈ I, J must be even closer to a singular matrix J_0_, i.e. J – J_0_ ≪ D – I. In this sense, the threshold D – I is asymptotically “large”. Understanding how a large threshold arises more generally outside this asymptotic regime and lowers as *N* increases remains an open problem, as do extending the present analysis to include the nonlocal interactions [48, 49] that arise for example in vegetation patterns [50] and extending previous work [16, 51] on the robustness of Turing patterns to *N* ≥ 3. The latter in particular may help to identify those chemical or biological pattern forming systems with *N* ≥ 3 in which the “true” Turing instabilities discussed here can be realized experimentally.

We thank N. Goldenfeld, A. Krause, and P K. Maini for discussions. This work was supported in part by a Nevile Research Fellowship from Magdalene College, Cambridge, and a Hooke Research Fellowship (P.A.H.), Established Career Fellowship EP/M017982/1 from the Engineering and Physical Sciences Research Council and Grant 7523 from the Marine Microbiology Initiative of the Gordon and Betty Moore Foundation (R.E.G.).

## Supplemental Material

This Supplemental Material is divided into five sections, which provide (i) details of calculations for *N* = 2, (ii) the derivation of the semianalytic approach for *N* = 3 and a discussion of its numerical implementation, (iii) an analysis of the statistics of the wavenumber at which a Turing instability first arises, (iv) a discussion of Turing instabilities with nondiffusing “slow” species, and (v) a proof of the asymptotic result claimed in the conclusion of our Letter.

### I. DETAILS OF CALCULATIONS FOR *N* = 2

#### A. Derivation of Eq. (3)

The form of the condition for Turing instability in Eq. (3) follows from that in Eq. (2.26) on page 85 of Vol. II of Ref. [S1] which, in our notation, reads

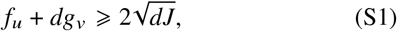

a quadratic in *d* = *d_u_ ≥ d_v_*. Hence

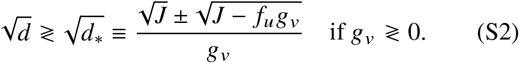

We notice that Eq. (S1) requires *f_u_* + *dg_v_* ≥ 0. Since *I*_1_ < 0, this implies that 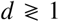 if 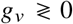. Hence 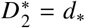 if *g_v_* > 0, but 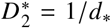 if *g_v_* < 0. Now, if 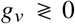, then 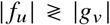 because *I*_1_ < 0 and *p* > 0. Equation (3) then follows, since

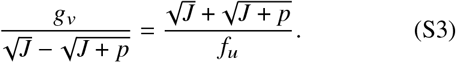

#### B. Derivation of Eq. (5)

Equation (3) shows that 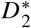 is continuous on *I*^4^, so attains its maximum value on that domain. Since *p* > 0 and *J* > 0, *q* ≡ – *f_v_g_u_* > 0, so that *J* + *p* = *q*. Now 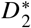 only depends on *f_v_, g_u_* through *q*, and, by direct computation from Eq. (3),

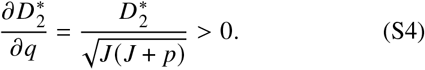

Hence 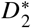 increases with *q*, so (*f_v_,g_u_*) = ±(*R, – R*) at the maximum.

Now assume that |*f_u_*| ≥ |*g_v_*|. Since *I*_1_ < 0 and |*f_u_*| ≥ |*g_v_*|, it follows that *f_u_* < 0 and *g_v_* > 0. Then

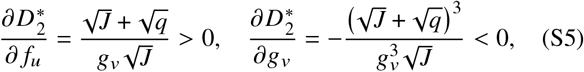

and so (*f_u_,g_v_*) = (1,-1) at the maximum. If |*f_u_*| ≤ |*g_v_*|, we similarly find that (*f_u_,g_v_*) = (−1,1) at the maximum. Substituting these values into Eq. (3) yields Eq. (5).

#### C. Calculation of 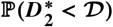 for 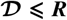

There are 48 ways of assigning values ± 1 and ±R to two of the entries *f_u_,f_v_,g_u_,g_v_* of J. Integrating the conditions for Turing instability of the remaining entries in each of these cases using Mathematica (Wolfram, Inc.) gives the area of parameter space in which a Turing instability arises,

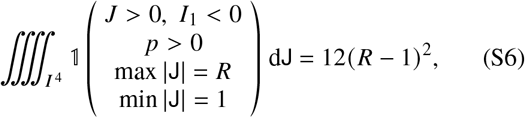

where we use the shorthand dJ = d *f_u_* d *f_v_* d*g_u_* d*g_v_*. To analyze the condition 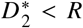, we note that the expression for 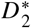 in Eq. (3) shows that we may swap *f_u_, g_v_* and *f_v_, g_u_*. Hence the 48 cases reduce to 4 cases (corresponding to the entries ±1 or ±*R* being on the the same or on different diagonals):

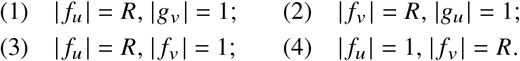

Moreover, since *q* > 0, we may take *f_v_* > 0 and *g_u_* < 0 without loss of generality. We now discuss these cases separately.

1. *I*_1_ < 0 implies *f_u_* = – *R*, *g_v_* = 1, and so

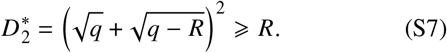
2. *f_u_g_v_* = – *R* since *q* > 0, so *J* = *f_u_g_v_* + *R*.
3. *f_u_* = – *R* because *I*_1_ < 0. Now *p,q* > 0, and so 0 < *J* = – *R*|*g_v_*| – |*g_u_*| < 0. This is a contradiction.
4. *f_u_* = 1 as *I*_1_ < 0. Since *g_v_* ≤ −1, it follows that

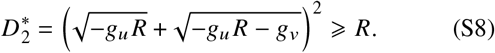

In this way, 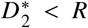 quantifies the diffusive threshold in a natural way. In particular, 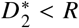 is only possible in case (2). Since *J* > 0, we require *f_u_ g_v_* + *R* > 0 in that case. Now *I*_1_ < 0 and *p* > 0, so 1 < *f_u_* < – *R*/*g_v_* or 1 < *g_v_* < – *R*/ *f_u_* depending on *f_u_* > 0, *g_v_* < 0 or *f_u_* < 0, *g_v_* > 0. Assume without loss of generality that |*f_u_* | ≥ |*g_v_*|. Then *f_u_* < 0, *g_v_* > 0 as *I*_1_ < 0. Moreover, using Eq. (3), 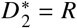 if and only if *g_v_* = 2 + *f_u_/R*. From Eqs. (S5), 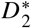 decreases as *g_v_* increases. Hence

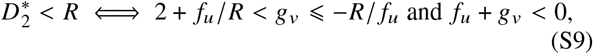

using the conditions derived previously. Note that – *R/f_u_* < *R* and 2 + *f_u_/R* > 1 for – *R < f_u_* < −1. If |*f_u_*| < |*g_v_*|, *f_u_,g_v_* are swapped in these conditions. Moreover, since *q* > 0, case (2) corresponds to 4 of the 48 cases. Hence we obtain, again using Mathematica,

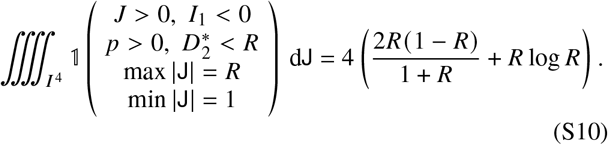

Equations (S6) and (S10) imply

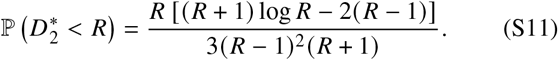

In particular, 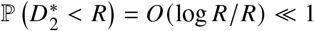 for *R* » 1. This statement expresses the existence of the diffusive bottleneck mathematically.

From a more physical point of view, as discussed in our Letter, it is more natural to consider the probability 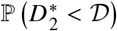, for some constant 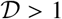. Since “small” values 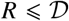 require fine-tuning of the reaction kinetics, we restrict to 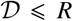, so that 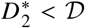 is only possible in case (2) above. We consider again the case *g_v_* > 0, *f_u_* < 0. Similarly to the derivation of conditions (S9), we find

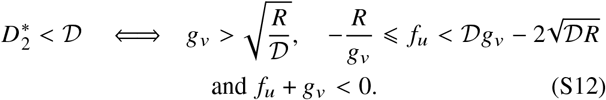

In particular,

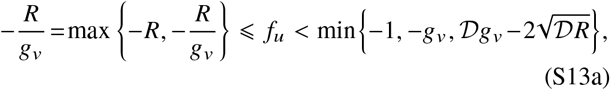

in which, since *g_v_* > 1,

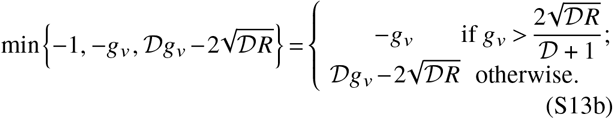

We notice that 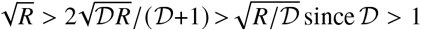, and also that 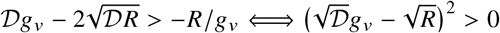, but 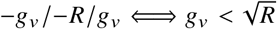. The area of parameter space described by conditions (S12) is therefore

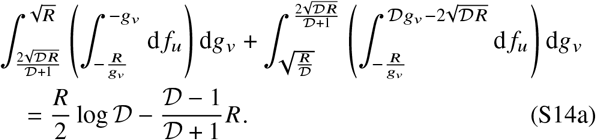

Hence [S2]

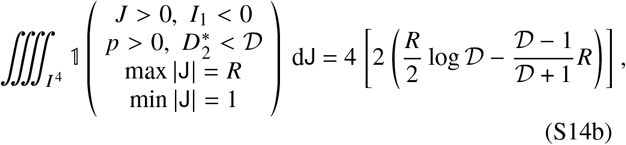

for 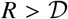, and, as above, we conclude that, for 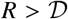,

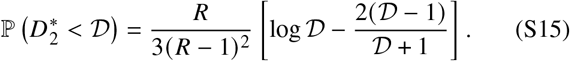

#### D. Nondimensionalization

We close by remarking on the (absence of) nondimensionalization of the reaction system. Indeed, up to rescaling time, one among *f_u_, f_v_, g_u_, g_v_* can be set equal to ±1. Moreover, one more parameter can be set equal to ±1 by rescaling *u, v* differently. However, if we made those choices, we could no longer sample from a fixed interval.

### II. SEMIANALYTIC METHOD FOR *N* = 3

#### A. Derivation of the semianalytic method

##### 1. Preliminary observations

Before deriving the semianalytic method, we need to make two preliminary observations.

First, the necessary and sufficient (Routh–Hurwitz) conditions for the homogeneous system to be stable include *I*_1_ ≡ tr J < 0 and *J* ≡ det J < 0 [S1]. By definition, J [*k*^2^) has one zero eigenvalue. The other two eigenvalues are either real or two complex conjugates *λ, λ*\*. In the second case, they are both stable (i.e. have negative real parts) since

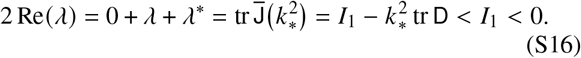

Hence Eqs. (7) are not unstable to an oscillatory (Turing– Hopf) instability at 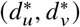, so, by minimality of 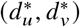, the system destabilizes to a Turing instability there.

Moreover, since 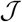, viewed as a polynomial in 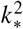, has leading coefficient – *d_u_ d_v_* and constant term 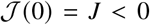, the double root *K*(*d_u_, d_v_*) varies continuously with *d_u_, d_v_* and cannot change sign on a branch of Δ(*d_u_, d_v_*) = 0 in the positive (*d_u_, d_v_*) quadrant.

##### 2. Reduction of problem (9) to polynomial equations

The discriminant of 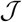, viewed as a polynomial in the two variables *d_u_, d_v_*, is

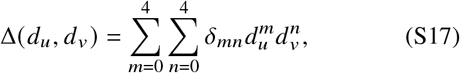

where *δ*_00_ = *δ*_10_ = *δ*_01_ = *δ*_34_ = *δ*_43_ = *δ*_44_ = 0 and (complicated) expressions for the 19 non-zero coefficients can be found in terms of the entries of J using Mathematica (Wolfram, Inc.).

The second remark above implies that, at a local minimum of *D*_3_ (*d_u_,d_v_*) on Δ(*d_u_,d_v_*) = 0, one of the following occurs:

a. Δ(*d_u_, d_v_*) = 0 is tangent to a contour of *D*_3_ (*d_u_, d_v_*);
b. Δ(*d_u_, d_v_*) intersects a vertex of a contour of *D*_3_ (*d_u_,d_v_*);
c. Δ(*d_u_, d_v_*) is singular.

The contours of *D*_3_ (*d_u_, d_v_*) are drawn in Fig. 2(a) of our Letter and show that tangency to a contour in case (i) requires

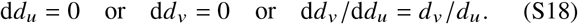

Since Δ(*d_u_,d_v_*) = 0, the chain rule reads

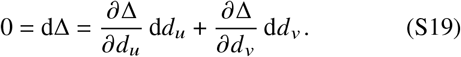

Hence there are two subcases:

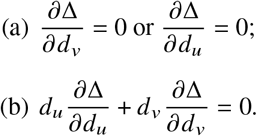

In subcase (a), Δ viewed as a polynomial in *d_v_* or *d_u_* has a double root, and so its discriminant [S3] must vanish. On removing zero roots, this discriminant of a discriminant is found to be a polynomial of degree 20 in *d_u_* or *d_v_*, respectively; complicated expressions for its coefficients in terms of the non-zero coefficients *δ_mn_* in Eq. (S17) are obtained using Mathematica. Similarly, in subcase (b), the resultant [S3] of Δ and *d_u_ ∂*Δ/*∂d_u_* + *d_v_ ∂*Δ/*∂d_v_*, viewed as polynomials in *d_u_* or *d_v_* must vanish. This resultant is another polynomial of degree 20 in *d_v_* or *d_u_*.

Next, in case (ii), *d_u_* = 1 or *d_v_* = 1 or *d_u_* = *d_v_* [Fig. 2(a)], which reduces *Δ* to three different polynomials in the single variable *d_v_, d_u_*, or *d = d_u_ = d_v_*, respectively. These polynomials have degree 6.

Finally, in case (iii), we note that, at a singular point, Δ = *∂*Δ/*∂d_u_* = *∂*Δ/*∂d_v_* = 0, and so we are back in case (i), subcase (a).

Thus, we have reduced finding candidates for local minima in (9) to solving polynomial equations: this defines our semianalytic approach. The global minimum is found among those local minima with *K*(*d_u_,d_v_*) > 0; in case (i), the roots only correspond to local minima if additionally *d_u_,d_v_* > 1 or *d_u_, d_v_* < 1 in subcase (a) and *d_u_* < 1 < *d_v_* or *d_v_* < 1 < *d_u_* in subcase (b) [Fig. 2(a)].

##### 3. Extension to binary systems with *N* > 3

For binary systems, the diagonal entries of D take two different values, *d*_1_, *d*_2_ only. Up to rescaling space, *d*_1_ = 1 and *d*_2_ = *d*, which turns the condition Δ(D) = 0 into 2^*N*-1^ – 1 different polynomial equations in the single variable *d*, corresponding to the different combinatorial ways of assigning diffusivities *d*_1_, *d*_2_ to the *N* species (in such a way that not all species have the same diffusivity). Determining the minimum value 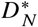 of *D_N_* = max {*d*, 1/*d*} for these binary systems is thus reduced, again, to solving polynomial equations.

The argument we used above to show that coexistence of Turing and Turing–Hopf instabilities is not possible for *N* = 3 does not, however, carry over to *N* > 3. Numerically, it turns out, however, that systems in which Turing and Turing–Hopf instabilities coexist are rare. We therefore treat these systems in the same way as we treat systems for which the numerics fail (as discussed below).

#### B. Numerical implementation

Implementing the semi-analytical approach for *N* = 3 and its extension to binary systems with 4 *≤ N ≤* 6 numerically takes some care as the coefficients of the polynomials that arise can range over many orders of magnitude. Our `python3` implementation therefore uses the `mpmath` library for variable precision arithmetic [S4].

To determine the positive real roots of the polynomials that arise in the semi-analytical approach, we complement the Durand–Kerner complex root finding implemented in the `mpmath` library [S4] with a test based on Sturm’s theorem [S3], to ensure that all positive real roots are found. Those systems in which root finding fails—either because the Durand–Kerner algorithm fails to converge or because it finds an incorrect number of positive real roots—are discarded, but included in error estimates where reported.

#### C. Numerical samples

Table S1 gives the number of random Turing unstable systems from which distributions, averages, and probabilities were estimated for each *R* ∈{2.5,5,7.5,10,12.5,15,17.5,20}.

For *N* = 3, we ran both a search for general, non-binary systems and a (larger but numerically less expensive) search for binary systems only. Since the first search only yielded binary global minima (as stated in our Letter), we used the results of the second, larger search for Figs. 3 and 4.

**TABLE S1.** Number of random Turing unstable systems used to estimate distributions, averages, and probabilities for the different values of *N*, and corresponding figures.

| $N$ | Type | $\max T^a$ | Reference<sup>b</sup> |
| --- | --- | --- | --- |
| $N = 2$ | non-binary | $10^7$ | Figs. 1, S1 |
| $N = 3$ | non-binary | $10^4$ | |
| $N = 3$ | binary | $10^5$ | Figs. 2, 4, S1 |
| $N = 4$ | binary | $10^5$ | Figs. 3, 4, S1 |
| $N = 5$ | binary | $2 \cdot 10^4$ | Figs. 3, 4, S1 |
| $N = 6$ | binary | $2 \cdot 10^3$ | Figs. 3, 4, S1 |

<sup>a</sup> Maximal number of Turing unstable systems.
<sup>b</sup> Figure (if any) in which results are shown.

### III. WAVENUMBER STATISTICS

In this Section, we discuss the wavenumber 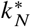 at which a Turing instability first arises at 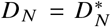. In particular, as discussed in our Letter, we must ask whether a Turing instability is “observable at the system size”. This observability requires the lengthscale 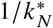 of the linear instability to be (a) smaller than the system *L* and (b) larger than *L/ℓ*, for some scale difference *ℓ* > 1. We are thus led to consider the probability 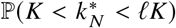, where *K* = 1/*L*.

It is instructive to start by considering the case *N* = 2. For the reaction-diffusion system in Eq. (1), a Turing instability arises for 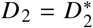 at a wavenumber 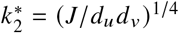 [S1]. We stress that this value depends on *d_u_, d_v_* not only through their ratio *d* = *d_u_/d_v_*. To absorb the dependence on the dimensional system scale, it is natural to consider

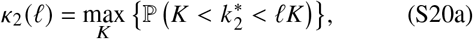

as the maximal probability of a Turing instability being observable at some inverse system scale *K* over a fixed scale difference Í. We denote by *K*_2_ (*ℓ*) the corresponding maximizing inverse system size.

For *N* > 2, we correspondingly ask: what is the probability of a Turing instability being observable at this inverse system size? We therefore define

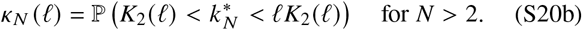

Figure S1 plots *κ_N_* (*ℓ*) against *N*, for fixed values of *R* and *ℓ*, but the qualitative behaviour is independent of *R* and *ℓ*. We notice that *κ_N_* (*ℓ*) increases slightly with *N*. If we restrict the analysis to those Turing unstable systems with 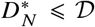, the probability is reduced somewhat for *N* > 2 compared to the case *N* = 2. This merely reflects the “fine-tuning problem”: the wavenumber is strongly constrained for those very rare systems that have a “small” diffusive threshold at *N* = 2. Moreover, about three quarters of the Turing instabilities at *N* > 2 do arise at physical wavenumbers, so we can extend the observations in Figs. 2(d) and 3(c) to note that random kinetic Jacobians are still more likely to be unstable to an observable Turing instability with small diffusive threshold for *N* > 2 than for *N* = 2.

**FIG. S1.**
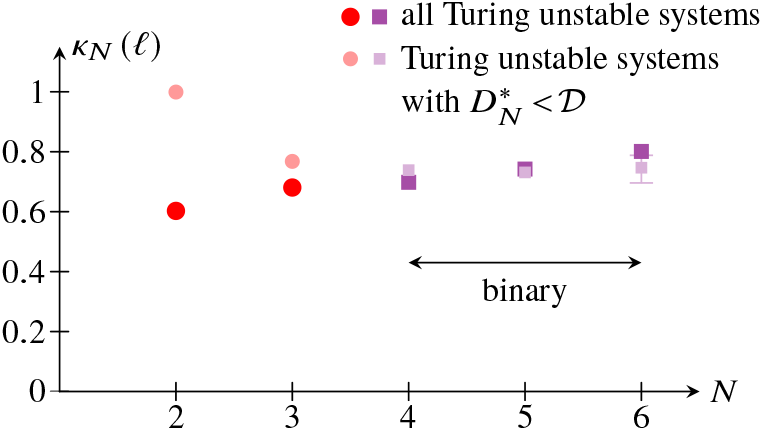
Wavenumber statistics. Probability *κ_N_* (*ℓ*) of a Turing instability being “observable” at a scale difference *ℓ* plotted against *N*; see text for further explanation. Larger markers: *w* (*ℓ*) estimated from all Turing unstable systems; smaller markers: (*ℓ*) estimated from only those Turing unstable systems with 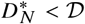. Parameter values: *R* = 10, *ℓ* = 10, 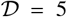. Asymmetric error bars again correspond to 95% confidence intervals larger than the plot markers, corrected for systems for which the numerics failed.

### IV. DIFFUSION OF “SLOW” SPECIES

In the notation of Eq. (12) of our Letter, Ref. [S5] shows that Turing instability at *d* = 0 requires J_22_ to be stable (i.e. all its eigenvalues to have negative real part): if it is not, instabilities arise at arbitrarily small and therefore unphysical lengthscales. In particular, det J_22_ ≠ 0, and so, using another result of Ref. [S5],

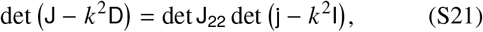

where 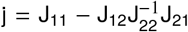. Hence a Turing instability occurs at *d* = 0 only if j has a positive real eigenvalue, as claimed in our Letter.

### V. THE ASYMPTOTIC DIFFUSIVE THRESHOLD

Let J = *O* (1) be a Turing unstable kinetic Jacobian, with an eigenvalue *λ* destabilising at nearly equal diffusivities, so that D = I + d with d = *o* (1). The following claim extends an argument of Ref. [S6]:

#### Claim

J *has a defective zero eigenspace.*

*Proof.* Because J – *k*^2^I has a stable eigenvalue *λ* – *k*^2^ and − *k*^2^d ≪ J – *k*^2^I, the corresponding eigenvalue of

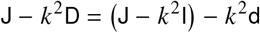

can only have positive real part if *λ* – *k*^2^ = *o* (1) i.e. if *λ* = *o* (1) and *k*^2^ = *o* (1) since Re(*λ*) < 0. Hence J and J – *k*^2^I have a zero eigenvalue at leading order. Additionally, the eigenvalue correction from – *k*^2^d = *o*(*k*^2^) must be *O*(*k*^2^) at least, which occurs iff the (leading-order) zero eigenspaces of J – *k*^2^I and J are defective [S7]; this final implication is discussed in more detail in Ref. [S8].

The generic case is therefore J = J_0_ + *O*(*ε*), where *ε* ≪ 1 and J_0_ has a defective double zero eigenvalue.

#### Claim

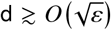; *in particular,* D – I » J – J_0_.

*Proof.* Since J_0_ has a defective double zero eigenvalue, J has two 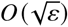 eigenvalues [S7], assumed to be stable (i.e. to have negative real parts). With *k* = *O*(*ε^κ^*), *d* = *O*(*ε^δ^*), destabilizing one of these requires, using the proof of the first claim above, – *k*^2^d ≳ *O*(*ε*) and 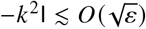, i.e. 2*κ + δ* ≤ 1 and *κ* > 1/4. Hence *δ* ≤ 1/2. This proves the claim.

## SUPPLEMENTAL CODE

The online Supplemental Material also includes excerpts from the `python3` code that we have written to implement the semianalytic approach for *N* > 3.

